# Withdrawal from repeated nicotine vapor exposure increases somatic signs of physical dependence, anxiety-like behavior, and brain reward thresholds in adult male rats

**DOI:** 10.1101/2022.01.08.475467

**Authors:** Michelle Martinez, Kevin Uribe, Valeria Garcia, Omar Lira, Felix Matos-Ocasio, Kenichiro Negishi, Arshad M. Khan, Laura E. O’Dell, Ian A. Mendez

**Author notes:** **Corresponding author:** Ian. A. Mendez, Ph.D., Assistant Professor, Department of Pharmaceutical Sciences, The University of Texas at El Paso School of Pharmacy, 500 W. University Drive, El Paso, TX 79902.

## Abstract

In recent years, there has been a dramatic increase in nicotine vapor consumption via electronic nicotine delivery systems (i.e., e-cigarettes), particularly in adolescents. While recent work has focused on the health effects of nicotine vapor exposure, its effects on the brain and behavior remain unclear. In this study, we assessed the effects that cessation from repeated nicotine vapor exposure had on behavioral and neuronal measures of withdrawal. For Experiment 1, fifty-six adult male rats were tested for plasma cotinine levels, somatic withdrawal signs, and anxiety-like behavior in the elevated plus maze, immediately following precipitated withdrawal from repeated exposure to 12 or 24 mg/mL nicotine vapor. In Experiment 2, twelve adult male rats were tested for intracranial self-stimulation (ICSS) across 14 days of exposure to 24 mg/mL nicotine vapor and across the 14 days immediately following nicotine exposure. Results revealed that plasma cotinine, somatic signs, anxiety-like behavior, and ICSS stimulation thresholds were all observed to be elevated during withdrawal in the 24 mg/mL nicotine group, when compared to vehicle controls (50/50 vegetable glycerin/propylene glycol). The data suggest that cessation from repeated nicotine vapor exposure using our preclinical model leads to nicotine dependence and withdrawal, and demonstrates that the vapor system described in these experiments is a viable pre-clinical model of e-cigarette use in humans. Further characterization of the mechanisms driving nicotine vapor abuse and dependence is needed to improve policies and educational campaigns related to e-cigarette use.

**Highlights:**

- A rodent model of nicotine e-cigarette vapor use was utilized to assess effects of cessation from repeated nicotine vapor exposure on behavioral and neuronal measures of drug withdrawal.
- Cessation of repeated nicotine vapor exposure resulted in increased plasma cotinine levels, somatic withdrawal signs, and anxiety-like behavior.
- Cessation of repeated nicotine vapor exposure resulted in elevations of ICSS reward threshold.
- Electrode implantations for ICSS were mapped by location and threshold to a standardized reference atlas of the rat brain to facilitate comparisons with the published literature.

## 1. Introduction

Addiction to drugs of abuse, such as nicotine, is a complex disease that is characterized by many components. Koob and Le Moal (2001) describe an addiction cycle that is comprised of three major components: preoccupation-anticipation, binge-intoxication, and withdrawal-negative affective states. Importantly, the authors note that different drugs and patterns of use can result in differences in the intensity of each component of the addiction cycle. One recent study investigated the abuse liability (i.e., tendency to use the drug excessively) of e-cigarettes against Food and Drug Administration approved nicotine inhalers and users’ own brand of traditional cigarettes. This study found that e-cigarette abuse liability was higher than that seen with nicotine inhalers, but lower than that observed with traditional cigarettes (Maloney et al., 2019). Additionally, comparable plasma nicotine levels following exposure to the highest nicotine concentration against traditional cigarettes suggests the potential for nicotine dependence with repeated e-cigarette use.

Increases in drug intake and behavioral measures of withdrawal, following repeated exposure to nicotine using intravenous self-administration and osmotic mini pumps has been well documented, particularly in rodents (Flores et al., 2020; O’Dell et al., 2007). A recently published study reports important initial steps for characterizing the effects of nicotine vapor on drug intake and withdrawal. In this study, researchers demonstrated that previous exposure to nicotine vapor during adolescence results in an increased propensity to self-administer nicotine later in adulthood through intravenous self-administration (Kallupi et al., 2019). The authors also demonstrate that while spontaneous withdrawal from nicotine vapor did not increase anxiety levels as indicated by the elevated plus maze, increases in somatic withdrawal signs were observed.

Research with rats has also shown that repeated nicotine exposure may also result in neuroadaptations, with aberrant dopamine signaling in the brain reward system being perhaps the most well-defined (Koob & Le Moal, 2005, 2008). The brain reward system can be studied in rodents by using Intracranial Self-Stimulation (ICSS) of the mesolimbic dopamine reward pathway (Markou & Koob, 1992) that projects from the *ventral tegmental area (Tsai, 1925)* (VTA) to the *accumbens nucleus (Ziehen, 1897–1901)* (ACB). Stimulating electrodes are aimed at specific structures of the brain reward pathway, including the *medial forebrain bundle* (*Edinger, 1893*) (mfb), a relatively larger section of the mesolimbic pathway (Carlezon & Chartoff, 2007). Electrical self-stimulations of the mfb are initiated through a response manipulandum and are frequently measured in amplitudes. Elevations in ICSS reward thresholds relative to baseline reflect decreases in reward sensitivity observed during negative psychological states, such as those seen with withdrawal (Markou & Koob, 1992). Studies have demonstrated increases in ICSS threshold during withdrawal from repeated nicotine injections (Fowler et al., 2013; Johnson et al., 2008). Identifying brain reward threshold changes during and after nicotine vapor exposure is an important step towards validating the pre-clinical vapor model of e-cigarette use and characterizing the effects nicotine vapor has on the brain reward system.

In this study, we investigated the effects that cessation of repeated nicotine vapor exposure had on biomarkers, behavioral measures, and neuronal mechanisms indicative of nicotine dependence. Specifically, Experiment 1 assessed the effects that cessation of repeated nicotine vapor exposure had on plasma cotinine levels, somatic withdrawal sings, and anxiety-like behavior, while Experiment 2 assessed the effects that repeated nicotine vapor exposure and cessation of repeated nicotine vapor exposure had on ICSS thresholds.

## 2. Methods

### 2.1 Nomenclature

In order to interrelate our ICSS data with those from other studies, we have opted to utilize the standardized neuroanatomical terms defined by Swanson (2015; 2018). These terms are listed in italics together with the associated citation that first uses the term as defined. If a definitive assignment of priority for the term was not possible, it was assigned by Swanson the citation “*(>1840)*”; that is, “defined sometime after the year 1840′′. Please refer to Swanson (2015, 2018) for further details regarding this standard nomenclature system.

### 2.2 Subjects

For Experiment 1, adult male Sprague-Dawley rats (7 weeks old upon arrival, n=56) were obtained from an outbred stock of animals (Envigo Inc., Indianapolis, IN). Experiment 2 used adult male Wistar rats (5 weeks old upon arrival, n=16) obtained from an outbred stock of animals (Envigo Inc., Indianapolis, IN). All animals were housed in a humidity-and temperature-controlled vivarium (22°C) with a 12-hr light/dark cycle (lights off at 6:00 AM and on at 6:00 PM) and *ad libitum* access to food and water. Animals were pair housed for Experiment 1, but individually housed throughout Experiment 2 to minimize electrode implant displacement. All described procedures were reviewed and approved by the Institutional Animal Care and Use Committee (IACUC) at the University of Texas of El Paso. Animals were allowed to acclimate to the vivarium for at least 1 week prior to any handling, training, or testing.

### 2.3 Apparatus

#### 2.3.1 Nicotine Vapor System

A Four Chamber Benchtop Passive E-vape Inhalation System (La Jolla Alcohol Research Inc., San Diego, CA) was used to deliver nicotine vapor to animals. The system is composed of four chambers large enough to house two rats per chamber (interior dimension of 14.5” L × 10.5” W × 9.0” H). Each chamber has two valve ports that are located on opposite walls and allow connection to vapor tubing. The output valve port was connected via PVC tubing to a small vacuum that created negative pressure in the chamber with an airflow at 0.6 L per minute. The vacuum outlet was connected to a Whatman HEPA filter (Millipore Sigma, Darmstadt, Germany), then onto a building exhaust system that safely removed the nicotine vapor from the chambers. The input valve port was connected via PVC tubing to a TFV4 mini-tank (4.9-volt, 65.0 W; Smok Inc, Shenzhen, China), where the nicotine e-liquid was heated by a 0.42 Ω atomizer coil. Each nicotine concentration (24 mg/mL, 12 mg/mL) and the vehicle control (propylene glycol/vegetable glycerin) was assigned its own tubing. Exposure chambers were thoroughly cleaned between exposures to avoid cross-contamination between experimental groups.

#### 2.3.2 Operant chambers

Operant chambers (12” L × 9.5” W × 11.5” H, Med-Associates, Inc., Fairfax, VT) used for ICSS were Plexiglas boxes, each with grid flooring, houselights, and a fixed-wheel manipulandum attached to the box wall. The wheel manipulandum is used to deliver electrical brain reward self-stimulations. A stimulator provides electrical stimulations that are sent through a commutator (2-channel, 305-plugs) located in each operant chamber. The commutator is attached to a commutator balance arm (0–20.3 cm, above chamber). A spring leash (5–100 cm, 2-channel; PTechnologies, Roanoke, VA) is screwed onto the commutator on one end and an electrode pedestal implanted into the animal on the other. A modified top permits the animal to move freely during training.

#### 2.3.3 Elevated plus maze

The elevated plus maze (EPM; Med-Associates, Inc., Fairfax, VT) was used to assess anxiety-like behavior, as a measure of withdrawal. The EPM consists of two closed arms and two open arms (10 cm W × 50 cm L). The closed arms had walls that were 20 cm in height and the maze was elevated 50 cm from the ground. All EPM testing was conducted under a red light, as differences in light exposure may influence behavior when using the elevated plus maze.

### 2.4 Drugs

For Experiment 1 a total of seven groups (n=8/group) were used and included: home cage (HC), vapor chamber (VC), 0 mg/mL vehicle control, 12 mg/mL nicotine vapor, 24 mg/mL nicotine vapor, 0.8 mg/kg nicotine injection, and saline injection. The HC group remained in their home cages for the entirety of the experiment, while the vapor chamber group was placed in vapor chambers without any vapor exposure. Flavorless nicotine e-liquids containing nicotine in its freebase form in a 50/50 vegetable glycerin/propylene glycol vehicle were used for this experiment. Nicotine e-liquids were purchased from a commercial vendor (Vapor Chef, VC Tobacco #13; Bristol, PA). Nicotine e-liquid concentrations of 24 mg/mL and 12 mg/mL, as well as a 0 mg/mL nicotine vehicle control were used, along with injections of nicotine ditartrate salt (0.8 mg/kg, s.c., expressed as a base) and the non-selective nicotinic receptor antagonist mecamylamine (3.0 mg/kg, s.c., salt, National Institute on Drug Abuse; Bethesda, MD). Both of these drugs were dissolved in 0.9% sterile saline and the pH of nicotine ditartrate salt was adjusted to 7.4 using a pH meter with chloride and hydroxide titration. For Experiment 2, a 0 mg/mL vehicle control (50/50 vegetable glycerin/propylene glycol) group and a 24 mg/mL nicotine vapor exposure group were used (n=6/group).

### 2.5 Experiment 1 Procedures

See **Figure 1** for timeline of Experiment 1 (Fig 1A) and Experiment 2 (Fig 1B) procedures.

**Figure 1.**
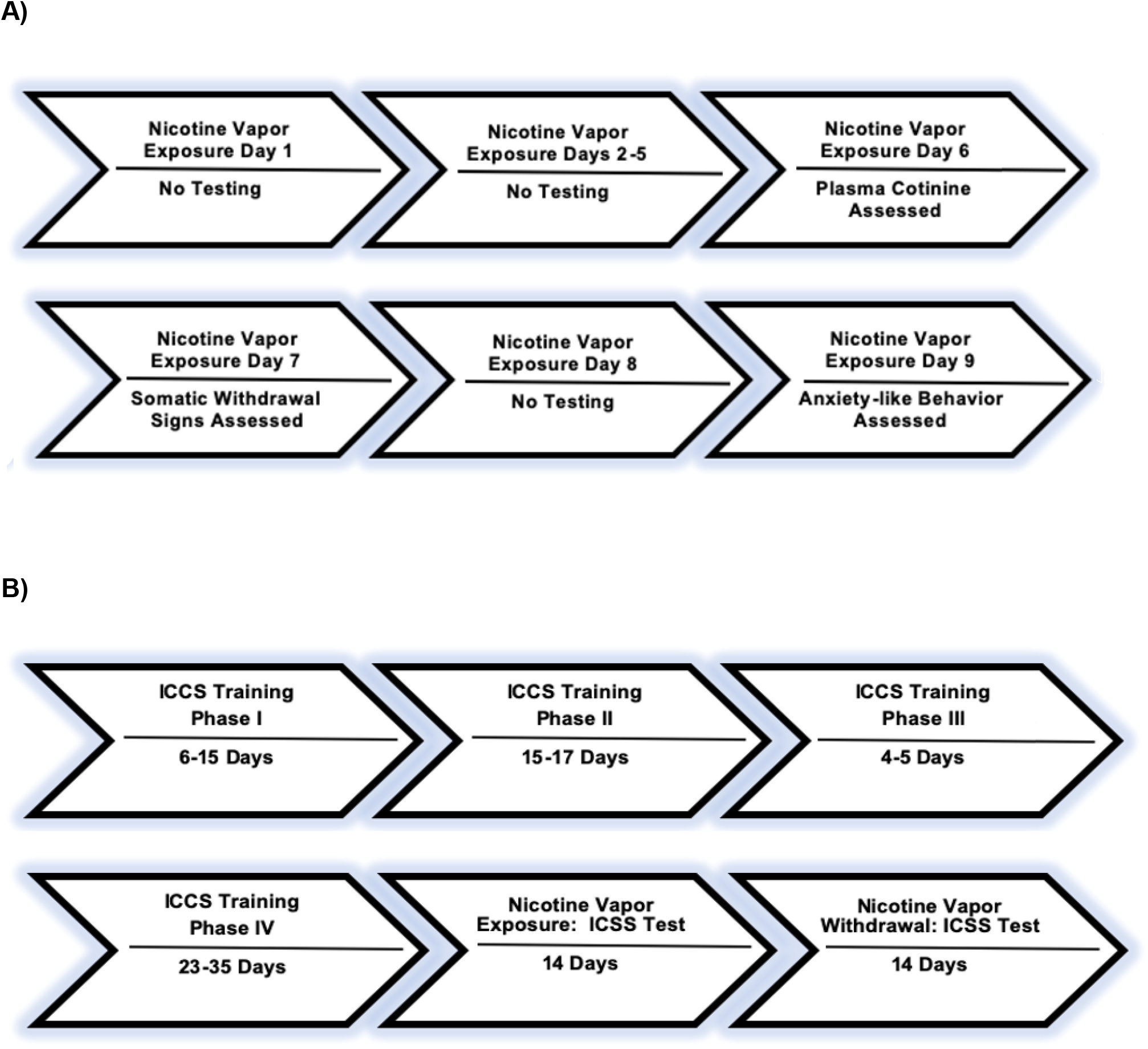
Experimental timeline for Experiment 1 (A) and Experiment 2 (B).

#### 2.5.1 Nicotine vapor exposure

For daily vapor exposures, animals with vapor exposure were placed in chambers with their respective cage mate. Pseudo-randomized assignment to vapor exposure groups was used due to the paired housing and exposure of the animals. The total time of nicotine vapor exposure sessions was 89 minutes for each nicotine concentration. Each session consisted of four cycles that lasts eighteen minutes and thirty seconds with an additional five-minute inter-exposure interval between each cycle. A 3-second nicotine puff is delivered into each chamber at the beginning of a cycle followed by a two-minute inter-puff interval, for a total of ten 3-second puffs every cycle. To model average daily nicotine vapor consumption in human e-cigarette users, a total forty puffs were delivered to the rats per daily nicotine vapor exposure sessions (Qasim et al., 2018). To help minimize nicotine contamination, lower nicotine concentration groups underwent vapor exposure before higher concentration groups during daily exposure sessions.

#### 2.5.2 Tail blood collection

To confirm systemic absorption of nicotine using our nicotine vapor inhalation system, tail blood (0.5 mL) was collected for each nicotine concentration on exposure day 6. This ten-minute procedure included brief anesthetization of each animal using 2.5–3% isoflurane gas in oxygen. Blood was collected using Eppendorf vials and centrifuged at (6900 r.p.m. × 4°C), before blood serum extractions were collected in separate vials and stored in a collection box placed in a –80°C freezer. Post-surgical topical analgesics were applied to the tail after sample collection. Cotinine detection was assessed utilizing an ELISA kit (Cal Biotech, Inc., El Cajon, CA), allowing for duplicate of the same sample. All standards, reagents, and substrate amounts were determined based on instructions provided in the ELISA kit.

#### 2.5.3 Somatic sign measures

On Day 7 of nicotine vapor exposure somatic signs were measured in all animals. Animals were placed in a Plexiglas box (30 cm × 29 cm) for assessment of somatic signs. Following acclimation to the box, rats were injected with mecamylamine (3.0 mg/kg, s.c.) and an additional ten-minute waiting period was required before precipitated withdrawal sign assessment. Number of somatic signs were recorded for ten minutes based on the following list of known indicators of withdrawal: blinks, yawns, teeth chatters, gasps, writhes, body shakes, headshakes, ptosis, and grooming (Flores et al., 2020; Harris et al., 2011).

#### 2.5.4 Elevated plus maze (EPM)

On Day 9 of vapor exposure, anxiety-like behavior, reflected by entries in the EPM, was assessed in all animals. Animals acclimated to the testing room in their home cage for ten minutes, followed by an injection of mecamylamine (3.0 mg/kg, s.c.; Harris et al., 2011). As with somatic sign assessment, a ten-minute waiting period following mecamylamine administration was required before placing animals in the middle of the EPM, facing an open arm. Assessment of behavior during precipitated withdrawal was based on closed versus open arm entries, across five minutes.

### 2.6 Experiment 2 Procedures

#### 2.6.1 Electrode implantation

Animals were anesthetized using isoflurane 2.5–5.0% gas in oxygen. Once anesthetized, rats were administered an intraperitoneal injection of saline (3.0 ml), a subcutaneous injection of flunixin (0.2 ml), and a subcutaneous injection of lidocaine on the scalp (0.1 ml). Animals were then positioned and secured in a stereotaxic frame (manufacturer, make, model), with a flat-skull position established by matching dorsal-ventral coordinate measures. Following screw placements, a +5.0-mm incisor bar elevation was applied, and a unilateral untwisted stainless-steel unipolar electrode (0.39 mm = ID, 0.71 mm = OD; P1 Technologies, Roanoke, VA) was implanted into one hemisphere (left and right hemisphere, counterbalanced) with the tip of the electrode targeting the mfb at the level of the *lateral hypothalamic area (Nissl, 1913)* (LHA), using the following stereotaxic coordinates: AP: –0.5 mm from Bregma; ML: ± 1.7 mm; DV: –9.3 mm from the skull surface; (Paxinos & Watson, 2007). After electrode placement, 2 or 3 screws were secured with acrylic cement glue to create a “cap” around the surgical site. A postsurgical antibiotic (i.e., Neosporin) was administered according to institutional regulations. Rats recovered from surgery for 5–7 days before any behavioral training was initiated.

#### 2.6.2 ICSS training

Animals began training in the discrete-trial current threshold ICSS task (Markou & Koob, 1992), which was delivered across four phases. In phase I of training, animals were tested individually in ICSS operant chambers. All animals started training with a default stimulation frequency of 100 Hz and 120 microamperes (μA), which were delivered freely following ¼ rotations of the response wheel. If low wheel-responding was observed in an animal after approximately 5 minutes in phase I or II training, 10–20 μA adjustments were made to the ICSS stimulation within sessions. Animals were trained in phase I until the criteria of 100 spins of the response wheel was achieved within 5 minutes. For phase II training, trials begin with a noncontingent stimulation (500 ms duration and 100 Hz), followed by a variable post-stimulation response window (7.5 s) during which delivery of a second stimulus was contingent upon a ¼ turn of the response wheel. Animals were required to spin the response wheel for 30 contingent electrical stimulations, at 5 different inter-trial intervals (ITI). ITIs were presented in ascending order (1, 3, 5, 10, and 15 sec), with one ITI assigned per day. Non-contingent and contingent stimulations were matched and daily sessions began with μA-stimulations that produced successful responding on the previous training day.

In phase III, we attempted to identify μA thresholds that produced maximal responding to non-contingent stimulations in individual rats. To determine these thresholds, μA threshold “blocks”, consisting of three trials per block, were automatically adjusted within a daily session by the MedPC program based on each animal’s responding to the non-contingent stimulus. Cathodal pulse stimulations (50–100 Hz) were delivered with a 500-ms train duration. When no response was detected for two out the three trials at a given μA-block, the stimulation was increased by 5 μA. When responding was detected for two out the three trials at a given μA-block the stimulation decreased 5 μA. In each daily session, two descending and two ascending μA-block series were presented, starting with a descending session and progressing in an alternating manner. The threshold for each of the four series was defined as the midpoint between the two μA-blocks immediately preceding a shift in series order. For phase III training day 1, the starting μA-setting was based on the last μA-setting successfully presented on the last day of phase II. On all subsequent phase III training days, the starting μA-setting was based on the average of the four identified series thresholds, plus 30 μA. If an animal maintained responding for 4–5 days on phase III, they were moved to phase IV. If responding was not maintained across 4–5 days on phase III, they were returned to phase II training.

Phase IV followed the same procedures used in phase III training. During phase IV, the average of the four alternating μA-block series was identified as the rat’s threshold for that day. This phase ended when all thresholds were stable, i.e., that there was <10% variability of amplitude increases and decreases across three training days. Response latencies were also identified and defined as the point between the start of the non-contingent stimulus and a ¼ turn response on the wheel (Chellian et al., 2021).

#### 2.6.3 Nicotine Vapor Exposure

Once rats achieved stability criteria for phase IV, they were pseudo-randomly assigned into vehicle control or 24 mg/mL nicotine vapor and underwent 14 consecutive days of vapor exposure immediately prior to ICSS testing. On day 1 of exposure, acclimation to vapor chambers was achieved by placing the rats in the chambers (without vapor) for 30 minutes before vapor exposure began. For daily vapor exposures, animals were placed in chambers with their respective cage mate (Experiment 1) or individually (Experiment 2). Pseudo-randomized assignment to vapor exposure groups was used due to the paired housing and exposure of the animals. The total time of nicotine vapor exposure sessions was 89 minutes. Each session consisted of four cycles that lasted eighteen minutes and thirty seconds with an additional five-minute inter-exposure interval between each cycle. A 3-second nicotine puff was delivered into each chamber at the beginning of a cycle followed by a two-minute inter-puff interval, for a total of ten 3-second puffs every cycle. To model average daily nicotine vapor consumption in human e-cigarette users, a total of forty puffs were delivered to the rats per daily nicotine vapor exposure sessions (Qasim et al., 2018). To help minimize nicotine contamination, groups exposed to nicotine vapor from lower concentrations of nicotine e-liquid underwent vapor exposure before those exposed to nicotine vapor from higher concentrations e-liquids during daily exposure sessions.

#### 2.6.4 ICSS threshold testing

Animals were tested in ICSS using procedures described for Phase IV training. Testing occurred immediately after each daily nicotine vapor exposure session and for an additional 14 days following nicotine vapor exposure cessation. This approach allowed us to assess the effects of acute and repeated nicotine vapor exposure, and spontaneous withdrawal from repeated nicotine vapor exposure, on ICSS brain reward thresholds.

#### 2.6.5 Histology

Whole brains were collected following decapitation on withdrawal day 14, following ICSS testing, for electrode placement verification. Brains were stored for 3–4 days in 4% *p*-formaldehyde (PFA) diluted using 1× PBS, then placed in a 30% sucrose/70% PFA solution for six days. Brains were cut into 40 μm-thick sections using a cryostat (Leica, CM1860) and tissues were mounted onto gelatin-coated glass slides. The mounted tissue was rinsed using distilled water, and then dehydrated in ascending concentrations of ethanol (50%, 70%, 95%, and 100%; 3 min each) for compatibility with subsequent immersion in a xylene bath (Thermo Fisher Scientific, Waltham, MA). Slide-mounted tissues were then gradually rehydrated through descending ethanol concentrations (100%, 95%, 70%, and 50%, 3 min each) and then stained with thionine (Cat #, Thermo Fisher Scientific, Waltham, MA). Tissues were again dehydrated through ascending ethanol concentrations and xylene for compatibility with DPX mounting medium (cat#, supplier), and then left to air-dry. DPX was also used to coverslip the slides, which were then left to dry overnight.

#### 2.6.6 Microscopy and Imaging

Tissue sections were observed with under bright-field illumination at × 5 magnification (Fluar objective, N.A. 0.25, FN 23 mm) using a AxioImager M.2 upright microscope (Carl Zeiss Corporation, Thornwood, NY) and photographed using an EXi Blue monochrome camera (Teledyne QImaging, Inc., Surrey, British Columbia) with Volocity software (Ver. 6.1.1; Quorum Technologies, Inc., Puslinch, Ontario). Native images were captured as stitched mosaics of the entire tissue section and exported as TIFF-formatted files for standardized mapping of the electrode placements.

#### 2.5.7 Standardized mapping of electrode placements

TIFF-formatted images were loaded into Adobe Illustrator (AI; Version CC 24.1.2; Adobe Systems, Inc., San Jose, CA). Structural boundaries were identified using the cytoarchitectural definitions outlined in *Brain Maps 4.0* (*BM4.0*; Swanson, 2018), and drawn in a separate layer using the *Pencil Tool* in AI. For a given electrode placement, boundary assignments were prepared for thionine-stained tissue sections containing the deepest detectable necrotic fields and for the sections flanking them. Atlas-level assignments were performed on the basis of plane-of-section analysis and comparison with flanking sections. Electrode placement locations were defined as the ventral tips of the necrotic fields and were plotted on *BM4.0* digital atlas templates.

### 2.7 Statistics

For Experiment 1, Statistical analyses included a one-way ANOVA with LSD post-hoc tests for cotinine levels, somatic withdrawal signs, and open arm entries. Bonferroni post-hoc tests were also conducted for all analyses and significant LSD and t-tests that were also found to be significant with Bonferroni are denoted with an asterisk (*). For Experiment 2, statistical analyses (IBM SPSS, Chicago) used included mixed-model ANOVAs with t-tests post-hoc analysis for response latencies, threshold μA-settings, and threshold percent changes from baseline (average of the last three days of Phase IV training thresholds for nicotine vapor exposure and average of the last three days of nicotine vapor exposure thresholds for nicotine vapor withdrawal).

## 3. Results

### 3.1 Experiment 1

#### 3.1.1 Blood plasma cotinine levels

Figure 2A illustrates blood plasma cotinine levels from tail vein blood that was extracted from all groups following 6 days of nicotine vapor exposure. One-way ANOVA revealed a main effect of group [*F*_(6,49)_=157.58; *p*≤0.001]. Post-hoc analyses show that rats in the 12 mg/mL and 24 mg/mL nicotine vapor groups expressed higher cotinine levels than HC, VC, 0 mg/mL, and saline injection groups (*p*_*s*_≤0.001*). Similarly, post-hoc tests revealed significantly higher cotinine levels in the 0.8 mg/kg nicotine injection group when compared to the saline injection group (*p*≤0.001*). When comparing groups exposed to nicotine vapor exposure to group exposed to nicotine injections, post-hoc analyses show that the 12 mg/mL group had significantly lower cotinine levels (*p*≤0.001*), while the 24 mg/mL group had higher cotinine levels, than the 0.8 mg/kg nicotine injection group (*p*≤0.001*). Finally, a comparison of cotinine levels between the 12 mg/mL nicotine vapor group and 24 mg/mL nicotine vapor group showed that the 24 mg/mL group had significantly higher cotinine levels than the 12 mg/mL group (*p*≤0.001*), following 6 days of vapor exposure. In summary, higher nicotine concentration exposure resulted in increased levels of cotinine when compared to animals in control groups.

**Figure 2.**
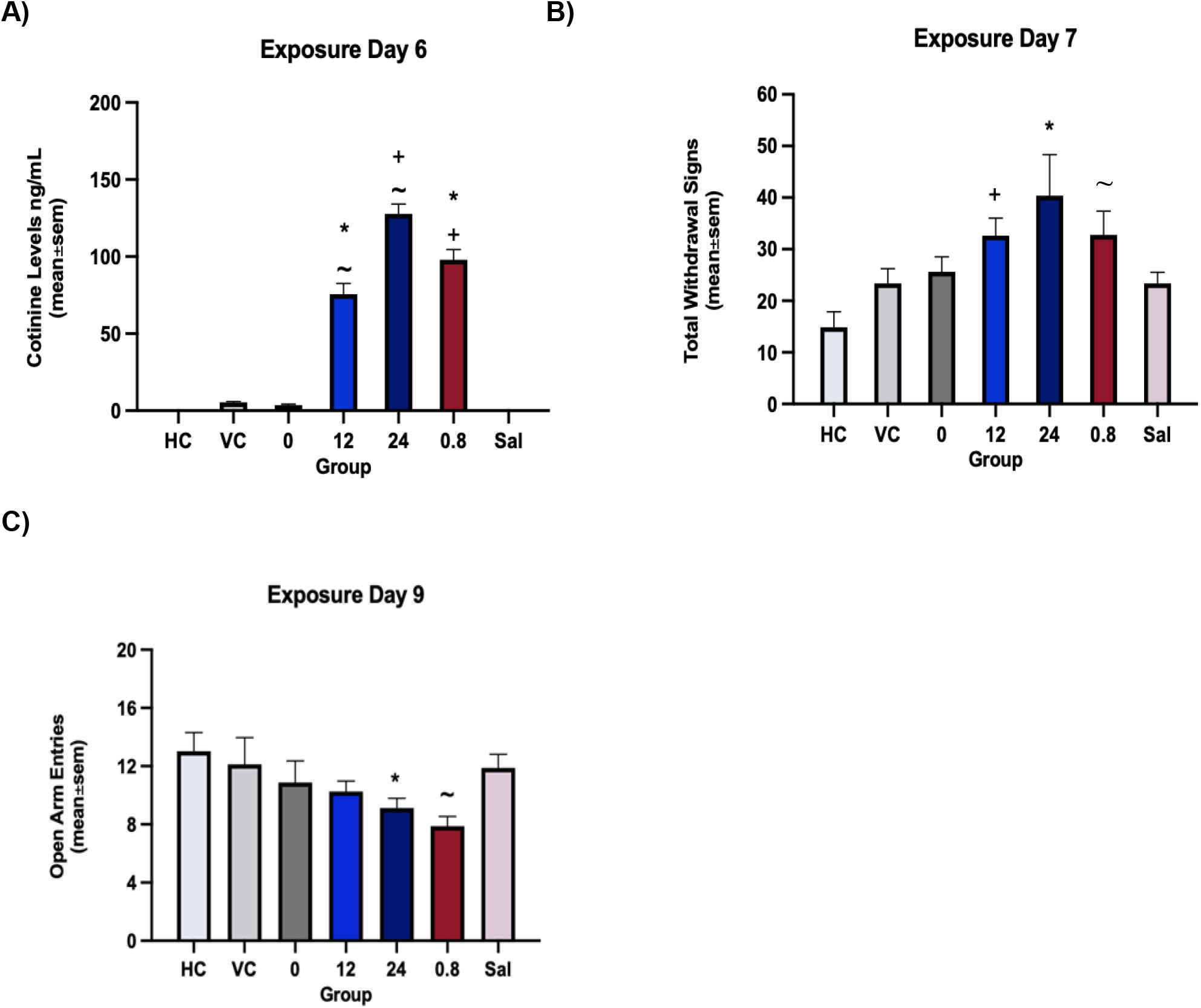
Blood plasma cotinine, somatic signs, and anxiety-like behavior following repeated nicotine vapor exposure. Animals exposed to nicotine vapor displayed higher cotinine levels (A), more physical withdrawal signs (B), and decreased open arm entries in the elevated plus maze (C). Asterisks (*) indicate significant difference from HC, VC, and 0 vapor control groups, plus sign (+) indicates significant difference from HC control group only, and tilde (∼) indicates significant difference from Sal control group. Critical *p*-value is 0.05.

#### 3.1.2 Somatic signs of withdrawal

Figure 2B illustrates somatic signs that were assessed in all groups on day 7 of nicotine vapor exposure, following mecamylamine administration. A one-way ANOVA revealed a main effect of exposure group [*F*(_6,49)_=3.84; *p*≤0.003]. Post-hoc analyses revealed that the 12 mg/mL and 24 mg/mL nicotine vapor group displayed more somatic signs when compared to the HC group (*p*≤0.005, *p*≤0.001*, respectively). The 24 mg/mL nicotine vapor group was also found to display more somatic signs than saline injection, VC, and 0 mg/mL vehicle control groups (*p*≤0.05). These findings demonstrate that increases in somatic withdrawal signs were displayed in animals exposed to higher nicotine concentrations when compared to controls.

#### 3.1.3 Elevated plus maze open arm entries

Figure 2C illustrates anxiety-like behavior that was assessed in all groups on day 9 of nicotine vapor exposure, following mecamylamine administration. A one-way ANOVA revealed a main effect of treatment group [*F*_(6,49)_=2.38; *p*≤0.05]. Post-hoc analysis revealed that the 24 mg/mL nicotine vapor group had fewer open arm entries than the HC group (*p*≤0.05) and that the 0.8 mg/kg nicotine injection group had fewer open arm entries when compared to the saline injection group (*p*≤0.05). Results in the EPM test show that exposure to higher nicotine concentrations resulted in fewer open arm entries in the EPM when compared to the control group.

### 3.2 Experiment 2

#### 3.2.1 ICSS Thresholds

Figure 3 illustrates ICSS threshold measures as μA values during the 14 days of nicotine vapor exposure, as well as the 14 days immediately following cessation of nicotine vapor exposure. Two rats from each treatment group were removed from the study, due to a failure to complete ICSS training or electrode dislodgement during the training, resulting in a final total of six rats per group. A mixed-model ANOVA comparing nicotine vapor exposure groups across the 14 days of treatment revealed a main effect of time [*F*_(13,126)_=1.84; *p* = 0.05], such that all rats exhibited a slight increase in threshold across the 14 days of treatment (Fig 3A). No main effect or interaction was observed for treatment group during nicotine vapor exposure days. Likewise, a mixed-model ANOVA comparing nicotine vapor exposure groups across the 14 days immediately following cessation of nicotine vapor exposure revealed no main effect or interaction for either time or treatment group (Fig 3B). Overall, thresholds between vehicle control and 24mg/mL nicotine groups, were comparable immediately following daily exposure to vapor. During withdrawal days thresholds between vehicle control and 24 mg/mL again remained similar.

**Figure 3.**
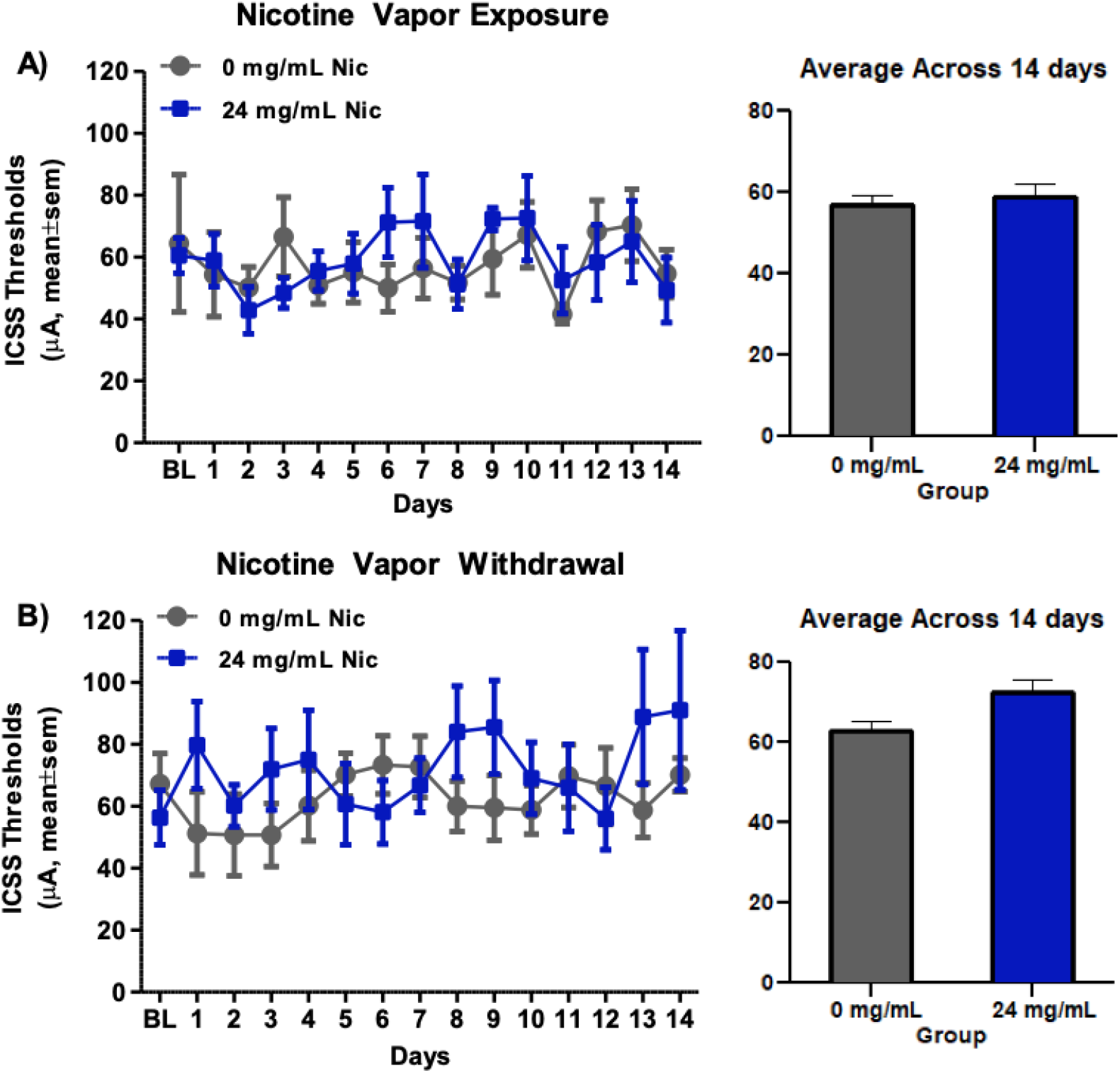
ICSS thresholds in microamperes during nicotine vapor exposure and withdrawal. No significant differences in reward threshold microamperes were seen across 14 days of vapor exposure (A) or across 14 days of withdrawal (B) when comparing animals exposed to 24 mg/mL nicotine vapor to animals exposed to vehicle control. Critical *p*-value is 0.05.

#### 3.2.2 ICSS Thresholds Percent Change

Figure 4 illustrates ICSS thresholds measures as percent change from baseline during the 14 days of nicotine vapor exposure, as well as the 14 days immediately following cessation of nicotine vapor exposure. A mixed-model ANOVA comparing nicotine vapor exposure groups across the 14 days of treatment revealed no main effect or interaction for time or treatment group (Fig 4A). Mixed-model ANOVA comparing nicotine vapor exposure groups across the 14 days immediately following cessation of nicotine vapor exposure revealed a main effect of group [*F*_(1,10)_=11.01; *p* = 0.01]. No main effect or interaction was observed for time during days immediately following cessation of nicotine vapor exposure. A post-hoc analysis comparing the nicotine vapor treatment groups on each day immediately following cessation of nicotine vapor exposure revealed higher percent changes in the 24 mg/mL nicotine vapor exposure group, when compared to the vehicle control group, on days 1, 3, 8, 9, and 13 (*p*_*s*_ = 0.05, Fig 4B). In summary, thresholds expressed as percent change were comparable in the vehicle control and 24 mg/mL nicotine groups immediately following daily vapor exposure. Interestingly, thresholds during withdrawal days were significantly different between the vehicle control and 24 mg/mL nicotine groups, with the 24 mg/mL nicotine group displaying higher threshold percent changes when compared to the vehicle control group.

**Figure 4.**
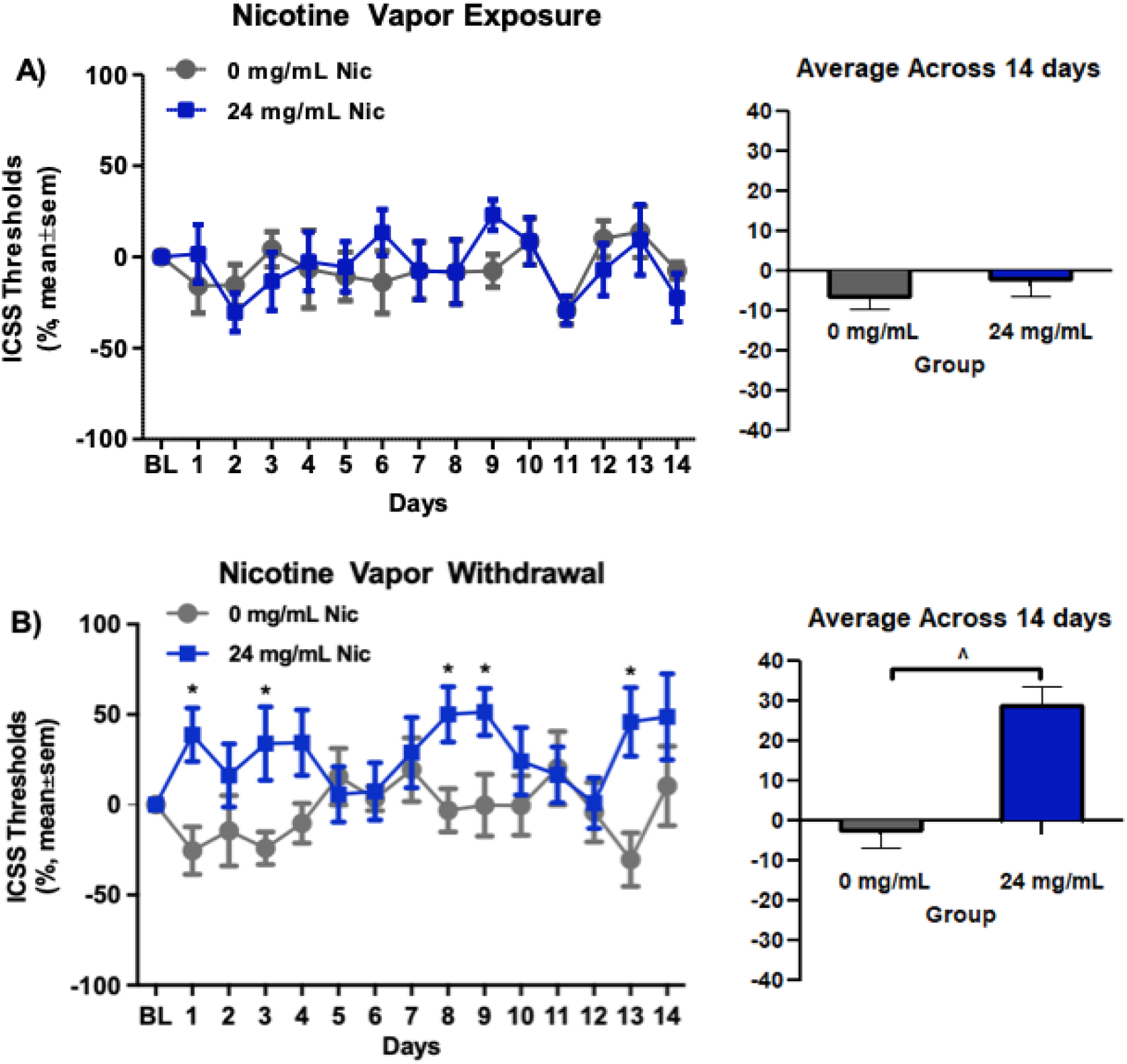
ICSS thresholds as percent change from baseline during nicotine vapor exposure and withdrawal. A) No significant differences were seen across 14 days of vapor exposure. B) Across 14 days of withdrawal the 24 mg/mL nicotine vapor group displayed a significantly higher threshold percent change from baseline than the vehicle control. The caret (^) indicates a significant main effect of treatment group across 14 days nicotine vapor withdrawal and an asterisk (*) indicates a significant difference between treatment groups on given day. Critical *p*- value is 0.05.

#### 3.2.3 Response latencies

Figure 5 illustrates ICSS response latencies during the 14 days of nicotine vapor exposure, as well as the 14 days immediately following cessation of nicotine vapor exposure. A mixed-model ANOVA comparing nicotine vapor exposure groups across the 14 days of nicotine vapor exposure revealed no main effect or interaction for time or treatment group (Fig 5A). Similarly, a mixed-model ANOVA comparing nicotine vapor exposure groups across the 14 days immediately following cessation of nicotine vapor exposure revealed no main effect or interaction for time or treatment group (Fig 5B). These findings suggest that exposure to 24 mg/mL nicotine does not significantly increase ICSS response latencies when compared to the vehicle control.

**Figure 5.**
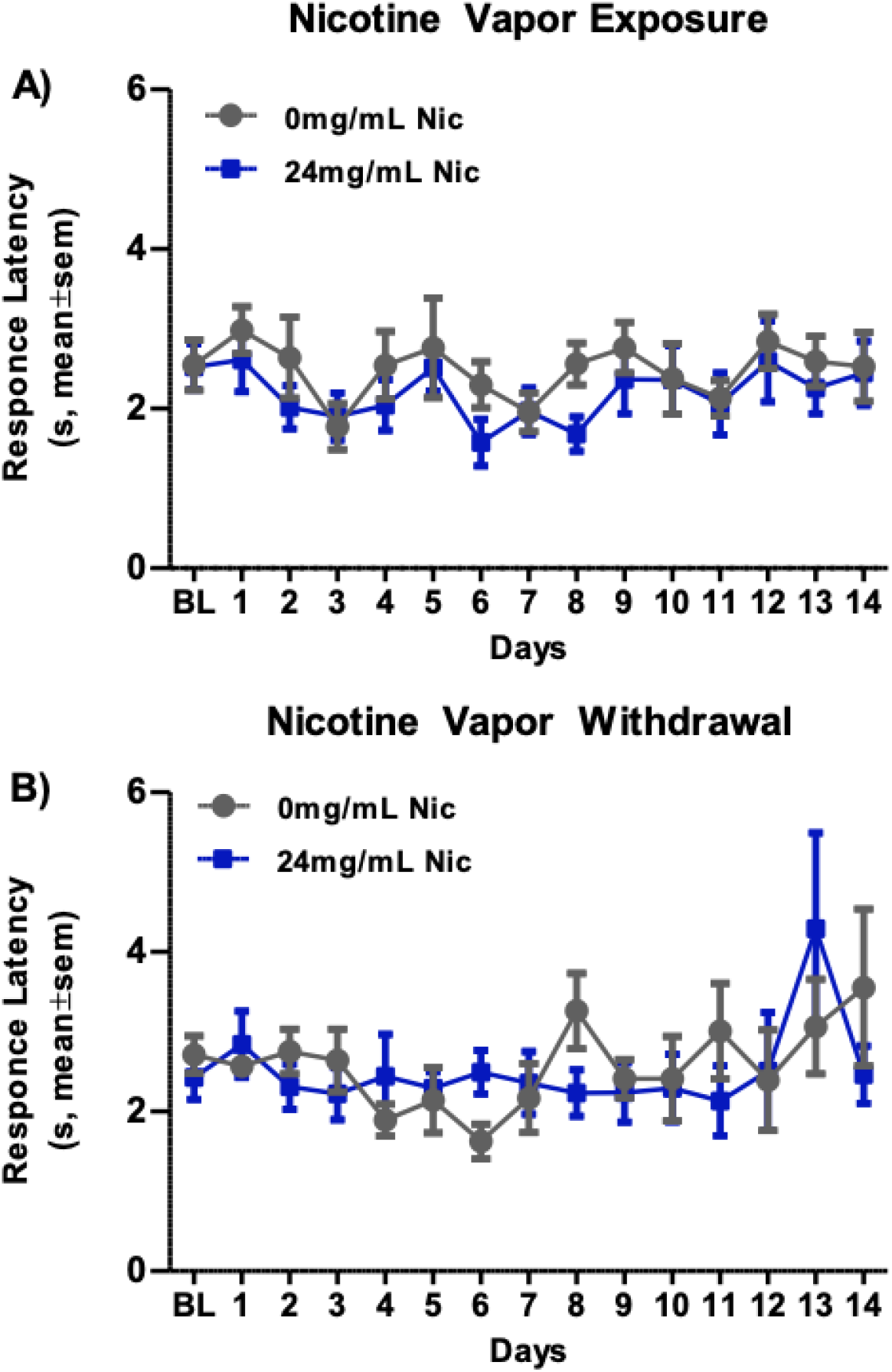
ICSS response latencies in seconds during nicotine vapor exposure and withdrawal. No significant differences in response latencies were observed across 14 days of vapor exposure (A) or across 14 days of withdrawal (B) when comparing animals exposed to 24 mg/mL nicotine vapor to animals exposed to vehicle control. Critical *p*-value is 0.05.

#### 3.2.4 Electrode placement verification

Figure 6 illustrates ICSS electrode placement in the mfb for all rats, as plotted onto vector-formatted templates of the *Brain Maps 4.0* rat brain reference atlas (Swanson & L.W., 2018). The histological analysis revealed that of the 12 rats completing ICSS training and test, 7 rats had placement in the mfb, with three rats having electrode placements just outside of the mfb (two from the 0 mg/mL vehicle control group and one from the 24 mg/mL vapor group) and two rats did not have confirmed electrode placements (one from the 0 mg/mL vehicle control group and one from the 24 mg/mL vapor group).

**Figure 6.**
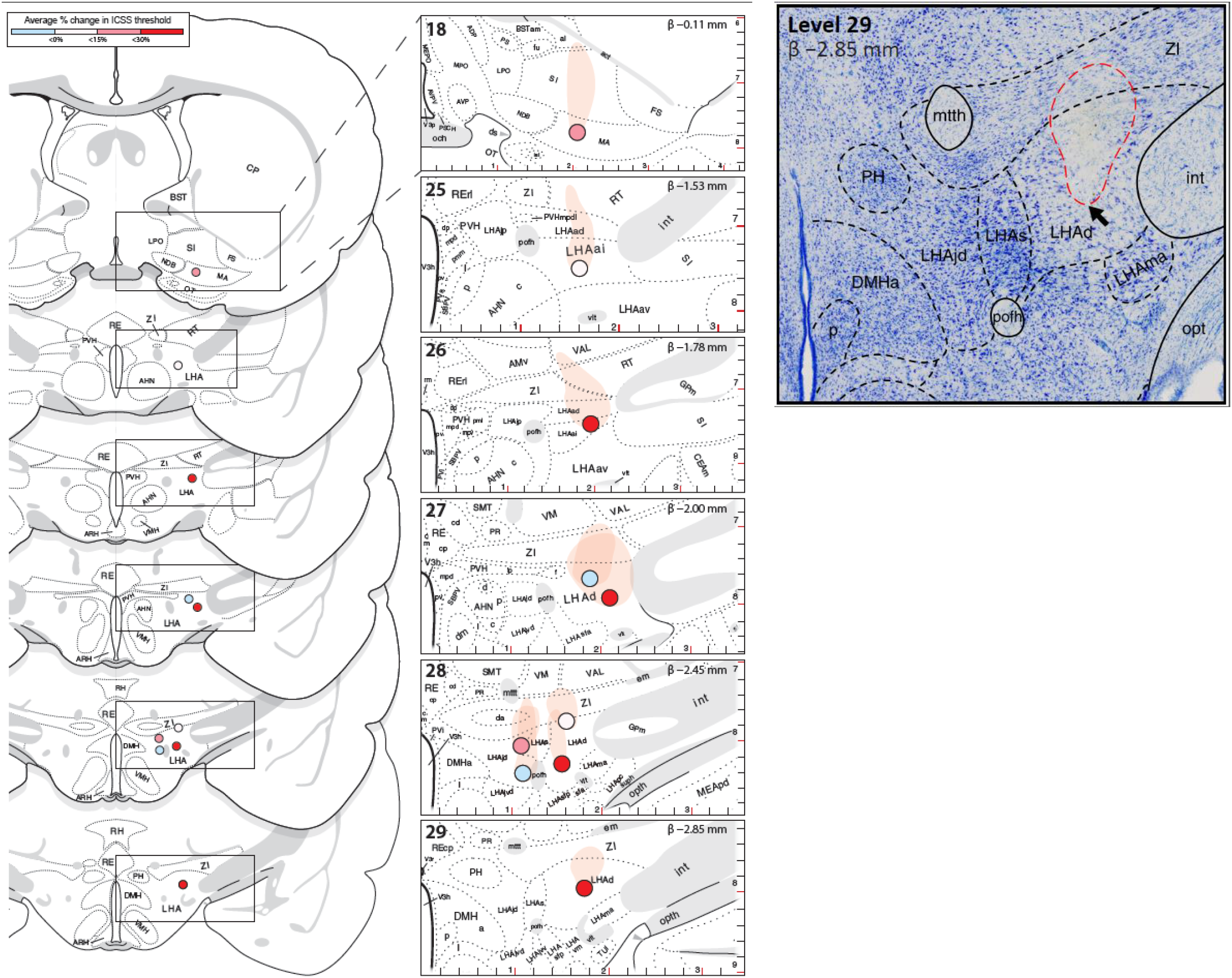
Mapped electrode locations in the *medial forebrain bundle (mfb; Edinger & L*., *1893)*, represented at two levels of resolution. A) The locations are denoted as dots and colored based on their average % changes in ICSS threshold. (*Left*) Wide view of ICSS probe locations across a 3-mm range of the mfb. (*Right*) Detailed view showing sub-regional architecture and inferred stereotaxic coordinates. The numbers on the top-left of each panel denote *BM4.0* atlas levels and those in the top-right indicate the corresponding Bregma coordinates (inferred for a flat-skull position). Vertical and horizontal rulers embed dorsoventral and mediolateral coordinates, respectively (1 increment = 0.2 mm). B) Representative thionine-stained section of an ICSS subject showing the location of an electrode implantation. The cell-sparse necrotic field is marked (red-dashed outline) along with the mapped site of the electrode tip penetration (black arrow). The *BM4.0* atlas level (Swanson, 2018) and Bregma coordinate (both top-left) were inferred based on the observable cytoarchitecture, which was parceled according to *BM4.0* atlas nomenclature (see Materials & Methods). The scale bar marks 500 µm.

## 4. Discussion

### 4.1 Experiment 1

The findings from Experiment 1 demonstrate that the described novel nicotine vapor system creates significant withdrawal-like behavioral states and increases biomarkers of withdrawal in male rats, following cessation of repeated nicotine vapor exposure. Specifically, this model produced significant increases in plasma cotinine biomarkers, somatic withdrawal signs, anxiety-like behavior, and c-fos activation in the habenula following cessation of nicotine vapor exposure, similar to that observed with more well-established routes of nicotine administration in rodents.

A major nicotine metabolite, cotinine, has a longer half-life (1519 hours) compared to nicotine (2–3 hours), and other plasma nicotine biomarkers (Avila-Tang et al., 2013; Benowitz, 1996; Buccafusco & Terry, 2003). This characteristic makes cotinine detection reliable for assessment of nicotine vapor exposure. Findings show that exposure to nicotine vapor (12 and 24 mg/mL) did indeed increase cotinine levels relative to negative controls on day 6 of exposure. When comparing animals in the 24 mg/mL group to animals in the 12 mg/mL group, we found that the group with the higher concentration of nicotine also had higher levels of cotinine, as expected. Additionally, we found that when compared to the group receiving 0.8 mg/mL nicotine injections across treatment days, the 24 mg/mL vapor group had slightly higher cotinine levels, and the 12 mg/mL group had slightly lower cotinine levels. The 0.8 mg/kg dose was selected as a positive control as it is considered a moderate injection dose of nicotine that has been shown to produce cotinine levels in rats similar to those seen in in traditional cigarette smokers (Allen et al., 2019; Vieira-Brock et al., 2013). While the nicotine e-liquid concentrations used in our experiment produce cotinine levels that were slightly outside of the range produced by the 0.8 mg/kg injection, they did produce cotinine levels comparable to those observed when using other traditional doses and routes of administration (Craig et al., 2014; Shoaib & Stolerman, 1999; Torres et al., 2013), as well as cotinine levels observed in human e-cigarette users (Flouris et al., 2013; Marsot & Simon, 2016). Others have demonstrated that the use of passive nicotine vapor exposure in rodents also effectively models pharmacokinetics related to behavioral and metabolic changes in human e-cigarette users (Shao et al., 2019). However, one limitation to consider with rodent vapor exposure, is the limited control of total nicotine administered to the subjects. Systemic delivery of nicotine in animals via vapor inhalation can indeed fluctuate based on subject size and inhalation rate, as well as location within the vapor exposure chamber. Overall, the translational framework of the described rodent vapor inhalation system can uniquely contribute to the literature on nicotine addiction by providing a reliable and valid animal model of human e-cigarette consumption (Matta et al., 2007).

Increases in physical withdrawal and anxiety-like behavior following cessation of nicotine exposure have been observed across animal species. Somatic behavioral signs of withdrawal have been thoroughly characterized in rodents (Malin et al., 1992), while decreases of open arm entries in an elevated plus maze are commonly assessed as a measure of anxiety-like behavior (Walf & Frye, 2007). Our behavioral procedures characterized both somatic signs and open arm entries in an elevated plus maze during precipitated withdrawal from repeated nicotine vapor exposure. The results revealed that rats exposed to 24 mg/mL nicotine vapor yielded more somatic withdrawal signs (e.g., blinks, yawns, writhes etc.) relative to those in all control groups, while rats exposed to 12 mg/mL nicotine vapor yielded more somatic withdrawal signs relative to only those in the home cage control group. Similar evidence suggests that repeated exposure to nicotine using more traditional routes of administration also leads to increases in somatic withdrawal signs (Skjei & Markou, 2003).

Regarding entries to the elevated plus maze open arms, analyses showed that animals receiving 24 mg/mL nicotine vapor had significantly fewer open arm entries than animals that were left in their home cages throughout the experiment. Decreases in open arm entries on an elevated plus maze have previously been observed in mice exposed to other concentration of e-cigarette vapor or to cigarette smoke (Ponzoni et al., 2015). Interestingly, a more recent study also showed decreases in open arm entries following nicotine vapor exposure in male and female rats, despite a 3-week abstinence period between last day of vapor exposure and testing (Smith et al., 2020), suggesting that nicotine induced anxiety-like behavior may be a long-lasting characteristic observed during withdrawal. To our surprise, our model produced only modest decreases in open arm entries when tested immediately following day 9 of nicotine vapor exposure. Studies have shown that females display increases in anxiety-like behavior compared to male rats on the EPM and these differences have been suggested to be mediated by ovarian hormones and stress responses (Flores et al., 2020; Torres et al., 2015). Similar results have also been observed in female mice following chronic nicotine exposure (Caldarone, King et al. 2008). Therefore, a limiting factor in the current study, possibly explaining the modest effect of anxiety-like behavior observed, was that we did not include female rats in our investigation. However, the EPM has shown to have face validity in its ability to measure anxiety in rodents, reflected by avoidance into open vs closed arms of the maze. In the EPM the animals must choose to enter into dark protected areas or into a lit open vulnerable space. Since rats are nocturnal animals they naturally prefer the darker protected areas. Other behaviors displayed in open arms such as freezing, immobility and defecation are anxiety-related and increased in open arms compared to closed arms (Pellow et al., 1985; Walf and Frye, 2007).

### 4.2 Experiment 2

The findings from Experiment 2 demonstrate brain reward changes following cessation of repeated nicotine vapor exposure, through relative increases in brain reward stimulation thresholds. Specifically, increases in ICSS brain reward thresholds were observed following cessation of repeated nicotine vapor exposure. While previous research investigating the effects of nicotine exposure on physical withdrawal and brain reward function have administered nicotine using injections, minipumps, infusions, and tobacco smoke exposure, the current study examine, for the first time, the effects of nicotine vapor exposure on physical withdrawal and reward function. Our analysis identified no significant changes in threshold currents or threshold percent change from baseline during repeated nicotine vapor exposure. The findings of the current study, however, are not entirely surprising, as previous literature examining ICSS thresholds during nicotine exposure have produced mixed findings (Chellian et al., 2021; Epping-Jordan et al., 1998; Harrison et al., 2002; Huston-Lyons & Kornetsky, 1992). The notion of variability in ICSS reward thresholds during nicotine exposure may be due to differences in sample size per group, differences in methodology used between experiments, and the resulting variation in pharmacokinetics. For example, differences in the effects of nicotine delivery via osmotic minipumps on ICSS have been attributed to differences in minipump infusion rates (Epping-Jordan et al., 1998; Xue et al., 2020). For our experiment, a larger sample size per group could have helped reduce variability in this task.

Unlike previous work examining ICSS thresholds during nicotine exposure, most previous studies using ICSS have shown elevations in reward thresholds in rats during nicotine withdrawal, regardless of administration method (Kenny et al., 2003; Muelken et al., 2015; Tan et al., 2019; Watkins et al., 2000; although see Kenny and Markou 2006). Such effects have also been observed following cessation of other drugs of abuse that include, but are not limited to, depressants, opiates and stimulants, (Chartoff et al., 2012; Holtz et al., 2015; Schulteis et al., 1995). As with previous reports on the effects of nicotine withdrawal on ICSS, the current study revealed significant changes in thresholds during withdrawal from nicotine vapor exposure.

No significant changes in response latencies were observed during or after repeated nicotine vapor exposure, suggesting that repeated nicotine vapor exposure does not produce immediate or long-term increases in behavioral response time. The effects of nicotine on response latencies in ICSS have produced mixed findings. For example, one study found that increases in nicotine injection doses (0.125, 0.25, and 0.5 mg/kg) produced no effect in response latencies in ICSS (Harrison et al., 2002). Alternatively, another more recent study found that injection of nicotine significantly decreased response latencies (Xue et al., 2020). While increases in response latencies should be expected due to nicotine’s well defined psychomotor stimulant effect, significant increases in response latencies could represent performance effects resulting from nicotine’s stimulant effects, rather than effects of nicotine on brain reward processes.

Therefore, another potential limitation of the present study is that locomotor activity was not directly assessed during ICSS training or testing sessions. Assessment of activity during inter-trial intervals can help identify locomotor effects of nicotine that are outside of contingent reward responding (Schaefer & Michael, 1986). It is important to note, however, that mixed results on locomotor activity do exist and are likely due to the differences in rat strains, nicotine e-liquid concentrations, nicotine’s biphasic locomotor effect, and route and timeline of administration (Rauhut et al., 2008; Samaha et al., 2005; Schaefer & Michael, 1986). For example, repeated intravenous infusions of 30 μg/kg nicotine have been found to produce locomotor sensitization, while repeated peripheral injections of this same dose resulted in tolerance to nicotine’s locomotor stimulant effect, suggesting that route of administration plays a role in nicotine’s effects on activity (Lenoir et al., 2013).

### 4.3 Conclusions and Future Directions

The current study provides much needed data on the behavioral and neural effects of repeated nicotine vapor exposure. While these findings help validate a useful preclinical model of e-cigarette use, it also promotes the need for further investigation of nicotine vapor exposure on the brain and behavior. Future studies should investigate effects of e-cigarette vapor in vulnerable populations, such as females, which display enhanced anxiety-like behavior during nicotine withdrawal (Flores et al., 2020). Additionally, a recent study suggests that differences in intake patters of nicotine vapor between males and females may underlie observed differences in cotinine levels (Lallai et al., 2021). Future work should also include an assessment of the effects of nicotine vapor on adolescents, due to the dramatic increase in e-cigarette use in this population and differences in nicotine withdrawal observed between adolescent and adults (O’Dell et al., 2006; Xue et al., 2020). In addition to investigating the effects of nicotine vapor exposure in specific populations, future studies should continue to investigate the unique pharmacology of nicotine e-liquid and vaping behaviors, including how different concentrations of nicotine, intake patterns, chemical additives (e.g., salt), and chemical reaction products (e.g., nicotyrine) contribute to its effects on the brain and behavior (Gholap et al., 2020; Sleiman et al., 2016).

The recent increase in e-cigarette use observed in adolescents and young adults is driven by a shift in risk perception related to the negative health effects of e-cigarettes (Bandi et al., 2021; Gorukanti et al., 2017). The dramatic increase of e-cigarette use in these populations highlights the immediate need for research on the effects nicotine vapor has on the brain and behavior. This data is needed for the development of important educational campaigns and regulation policies (Collins et al., 2019). Additionally, while many traditional cigarette smokers seek smoking cessation through e-cigarettes, studies have not provided clear evidence on whether these systems are effective for smoking cessation compared to other nicotine replacement therapies (Pokhrel & Herzog, 2015; Villanti et al., 2018). A more thorough investigation of the marketing, perceptions, and health effects of these novel nicotine delivery systems will be required to better understand their long-term impact on society, and particularly, in vulnerable populations.

## Acknowledgements

The authors would like to thank Priscilla Giner, Yanabel Grant, Shelby Towers, and Ana Polar for their guidance and assistance with this project. Research performed by IM, LEO, MM, KU, FMO, OL, and VG was supported by the National Institute on Drug Abuse (grant numbers DA052119, DA021274). Research performed by AMK and KN is supported, in part, by the National Institute of General Medical Sciences (grant number GM127251). KN is an Eloise E. and Patrick Wieland Graduate Fellow.

